# Atlas of Transcription Factor Binding Sites from ENCODE DNase Hypersensitivity Data Across 27 Tissue Types

**DOI:** 10.1101/252023

**Authors:** Cory C. Funk, Alex M. Casella, Segun Jung, Matthew A. Richards, Alex Rodriguez, Paul Shannon, Rory Donovan-Maiye, Ben Heavner, Kyle Chard, Yukai Xiao, Gustavo Glusman, Nilufer Ertekin-Taner, Todd E. Golde, Arthur Toga, Leroy Hood, John D. Van Horn, Carl Kesselman, Ian Foster, Ravi Madduri, Nathan D. Price, Seth A. Ament

## Abstract

There is intense interest in mapping the tissue-specific binding sites of transcription factors in the human genome to reconstruct gene regulatory networks and predict functions for non-coding genetic variation. DNase-seq footprinting provides a means to predict genome-wide binding sites for hundreds of transcription factors (TFs) simultaneously. However, despite the public availability of DNase-seq data for hundreds of samples, there is neither a unified analytical workflow nor a publicly accessible database providing the locations of footprints across all available samples. Here, we implemented a workflow for uniform processing of footprints using two state-of-the-art footprinting algorithms: Wellington and HINT. Our workflow scans the footprints generated by these algorithms for 1,530 sequence motifs to predict binding sites for 1,515 human transcription factors. We applied our workflow to detect footprints in 192 DNase-seq experiments from ENCODE spanning 27 human tissues. This collection of footprints describes an expansive landscape of potential TF occupancy. At thresholds optimized through machine learning, we report high-quality footprints covering 9.8% of the human genome. These footprints were enriched for true positive TF binding sites as defined by ChIP-seq peaks, as well as for genetic variants associated with changes in gene expression. Integrating our footprint atlas with summary statistics from genome-wide association studies revealed that risk for neuropsychiatric traits was enriched specifically at highly-scoring footprints in human brain, while risk for immune traits was enriched specifically at highly-scoring footprints in human lymphoblasts. Our cloud-based workflow is available at github.com/globusgenomics/genomics-footprint and a database with all footprints and TF binding site predictions are publicly available at http://data.nemoarchive.org/other/grant/sament/sament/footprint_atlas.

## INTRODUCTION

Regulation of gene expression by transcription factors (TFs) forms the basis for tissue and cell-type differentiation, arising from a complex interplay between TFs and the chromatin architecture in gene regulatory regions [1, 2]. In humans, genetic perturbation of TF binding sites is thought to be an important mechanism by which single-nucleotide polymorphisms (SNPs) influence risk for human disease [3-5]. Thus, characterizing the cell type-specific occupancy of TFs at their genomic binding sites is a critical goal in genomics, providing insight both into networks of TFs and their cell type-specific target genes, as well as the causal mechanisms underlying risk for human disease [6-9] [10].

Mapping human gene regulation requires comprehensive resources of tissue- and cell type-specific TF binding sites. Major efforts over the past decade have produced vast quantities of public epigenomic data that have dramatically expanded the functional annotation of the human genome [11-13], yet our understanding of cell type-specific TF binding sites remains far from complete. Annotation of TF binding sites based solely on the locations of sequence motifs is imprecise, since only ∼1% of motif instances are occupied by a TF at any given time [1]. Similarly, information about the locations of promoters and enhancers lacks sufficient specificity, as many genetic variants in these regions do not impact gene expression [2]. TF occupancy can be ascertained with high sensitivity and specificity through chromatin immunoprecipitation followed by deep sequencing (ChIP-seq), in which an antibody specific to a TF is used to pull down genomic DNA fragments occupied by that TF in a given sample. However, high quality ChIP-seq data have been generated for only a minority of all human TFs and often used standard cell lines rather than disease-relevant human tissues.

Genomic footprinting is a higher-throughput approach that predicts TF genomic occupancy by combining information from open chromatin assays (such as DNase-seq) with information about the locations of sequence motifs recognized by the DNA binding domains of transcription factors. DNase-seq assays are predicated on accessibility of genomic DNA to DNase I, where regions of open chromatin are susceptible to cleavage by DNase I. Binding of TFs and other DNA-binding proteins can lead to a relative difference in the number of cleavage events in discrete regions along the genome, resulting in a footprint [14]. Computational algorithms have been developed to identify footprints from high-throughput DHS data, typically using one of two different strategies: 1) sliding window approaches where the relative number of DNase cleavage events are counted along a sliding window of the genome, agnostic to the absence or presence of a TF binding motif [15-19], and 2) approaches that begin with the known location for a TF binding motif and model the DNase cleavage patterns around it for all sites in the genome [20-24]. Validation of these approaches typically has involved comparison of the footprints for individual TFs to binding sites found by ChIP-seq. Notably, the computational identification of footprints from high-throughput data remains an area of active research, as existing algorithms detect genomic occupancy for only a subset of TFs. Moreover, due to redundancy in the sequence specificity of TFs, footprinting generally cannot distinguish which member of a TF family is occupying a footprint. Nonetheless, the accuracy and reproducibility of TF binding site predictions from footprinting analysis has begun to rival that of ChIP-seq, and DNase-seq footprinting has successfully been used to predict the binding sites for hundreds of TFs in a parallel approach.

One of the most important applications of comprehensive atlases of TF binding sites will be to functionally annotate genetic risk variants for human diseases. Many studies have shown that disease-associated SNPs are enriched in gene regulatory regions, including open chromatin regions identified through DNase-seq and ATAC-seq experiments [3, 4, 25, 26]. However, GWAS risk loci are defined by large sets of genetically correlated SNPs with similarly strong statistical associations to disease, of which only a subset are thought to be functional and causal for disease risk. It remains controversial how many of these causal SNPs disrupt gene regulation by altering the specific base pairs occupied by TFs vs. other mechanisms. Several studies have identified risk loci for traits such as obesity and schizophrenia in which causal variants appear to functionally alter binding sites for key transcription factors [6, 7, 9]. However, other studies question the generalizability of this insight and indicate that TF binding sites in existing databases do not fully predict causal variants [8]. One explanation for this discrepancy is that existing TF binding site databases do not include sufficient amounts of epigenomic data from disease-relevant tissues. Since the gene regulatory consequences of non-coding SNPs are likely to vary dramatically across tissues and cell types [6, 27], these existing databases may therefore miss context-specific effects of variants on TF occupancy. In addition, there is considerable variability in the sensitivity and specificity of footprinting algorithms, and it is unclear which approaches will be best suited for this task.

Here, we developed a comprehensive resource of genomic footprints across 27 human tissues, using data from 192 DNase-seq experiments from the Encyclopedia of DNA Elements (ENCODE). Prior to our work, there was no publicly available, scalable workflow utilizing these data for the purpose of producing footprints. These analyses revealed an expansive landscape of tissue-specific genomic occupancy for 1,530 TFs. We validated our database based on ChIP-seq and eQTLs, and we demonstrated that tissue-specific footprints are strongly and specifically enriched for disease-associated genetic variation. We have made our footprint database and the underlying cloud-based computational workflow available in a user-friendly and intuitive format (links available in STAR Methods) [28].

## RESULTS

### A comprehensive atlas of genomic footprints across human tissues

ENCODE-generated DNase-seq FASTQ files from 192 experiments in 27 tissues were downloaded from the ENCODE data portal (encodeproject.org). The tissue-specific genomic occupancy of 1,515 TFs was then predicted through genomic footprinting analyses using a workflow pictured in Figure 1A and detailed in Methods. First, sequence reads were aligned to GRCh38 using SNAP [29]. As the DNase-seq data consist of short reads, we generated two alignments: one using the default 20 bp seed length and another using a 16 bp seed length.We then identified regions of open chromatin in each of the 192 experiments using F-seq, followed by detection of footprints using both the HINT and Wellington algorithms. Footprints detected in each of the 192 experiments were then grouped by tissue, producing 27 tissue-specific footprint maps, with separate maps for each seed size and footprinting algorithm. In general, seed size had only a modest impact, with ∼70% of the footprints having complete overlap between the two seed sizes (Figure S1A). Also, we observed only a moderate relationship between the number of footprints found in a sample and the depth of sequencing (Figure S1B). Overall, HINT identified more footprints than Wellington.

**Figure 1.**
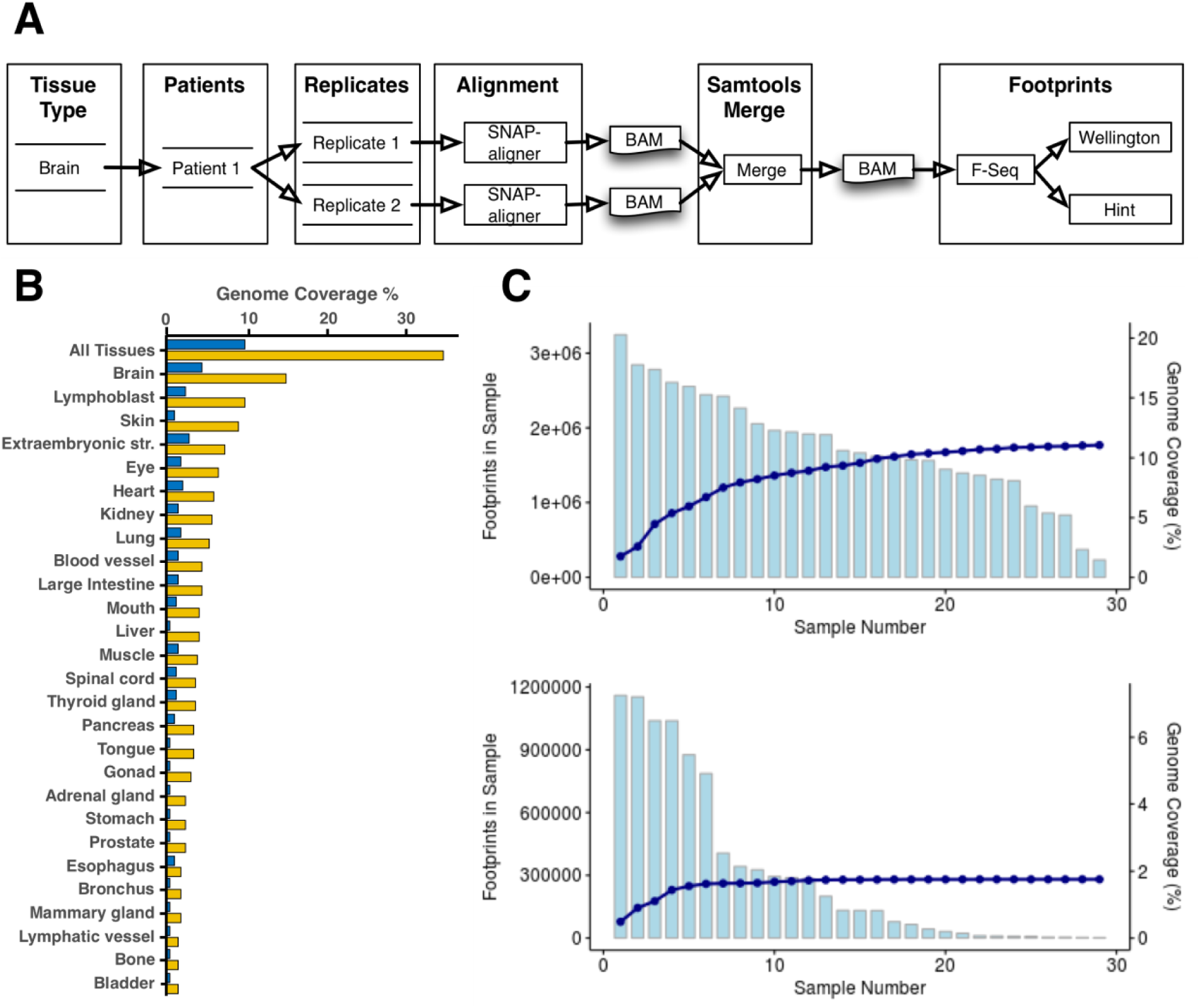
A) Footprints workflow overview - Each tissue type can have multiple quantity of patients and replicates. Each replicate is aligned using SNAP-aligner. All replicates for each patient are merged using Samtools. Finally, footprints for each BAM file are produced using Wellington and HINT and stored in a database. B) Percentage of the genome covered by the footprints for each tissue type and all tissues. Yellow is without filtering and dark blue is filtering HINT score > 200 and Wellington score < -27. (each method has its own scale and distribution) C) Footprints from the brain for HINT seed size 20 are ordered based on the number of footprints and summed. The light blue graphs represent the total number of footprints in each sample (top is without filtering on score; bottom is filtered as in panel B). The dark blue line represents the cumulative percentage of the genome covered.

Footprints from HINT and Wellington are identified without consideration of underlying motif sequence. Therefore, to predict which TFs occupy each footprint, we created a catalog of all genome-wide instances of 1,530 sequence motifs recognized by 1,515 TFs, using FIMO [30]. In addition to the motif-TF mappings provided by the aforementioned databases, we also expanded the motif-TF mappings to incorporate families of TFs with very similar DNA sequence specificity, using information from TFClass [31] (Tables S1, S2, Fig. S2). This resulted in ∼1.34 billion sequence-to-TF matches (p-value < 10^−4^) prior to intersection with footprints, spanning almost 80% of the genome. These motif instances were then intersected with the footprints from Wellington and HINT to produce an atlas of predicted TF occupancy in each tissue.

When considering all samples from all tissues, the most liberal thresholds resulted in 34% coverage of the genome being represented in the atlas for at least one tissue. The brain had the highest genome coverage at 14.9%, followed by skin (9.8%) and lymphoblast (8.9%). Urinary bladder had the lowest percentage of coverage at 1.1% (Fig. 1B). Sample size and sequencing depth were the main determinants for the number of tissue-specific footprints identified in our atlas. However, intrinsic biological differences in tissue complexity also influence the number of distinct footprint locations. For example, we found strong overlap in the footprint locations across the 46 experiments from skin (average pairwise Jaccard similarity index = 0.28), consistent with skin being a relatively homogeneous tissue. By contrast, the footprints detected in the 29 experiments from brain were less homogenous (average pairwise Jaccard similarity score = 0.16), which likely reflects the highly specialized and disparate cell types and cell type-specific gene regulation across brain regions. As a consequence, we identified more brain footprints than skin footprints, despite having 50% more skin samples.

An outstanding question is to what extent additional samples would add previously unseen footprints. To address this, using footprints derived from the HINT algorithm with seed length 20 (HINT20), we ordered the brain samples from most to fewest footprints and calculated the additional percentage of the genome covered by each sample (Figure 1C). The first sample contributed 3.25 million footprints spanning 1.75% of the genome, while the last sample added 235,000 novel footprints and 0.04% novel genome coverage. We repeated the same analysis using only high-quality footprints based on HINT and Wellington scores (see next section). As expected, this analysis revealed even greater overlap across samples, since many footprints detected in only a single sample are of low quality (Figure 1C, bottom). These results suggest that at least for well-sampled tissues such as brain, our atlas captures most detectable footprints.

### Validation and filtering of footprints with ChIP-seq and machine learning

Next, we sought to validate TF binding site predictions in our atlas and chose appropriate thresholds at which footprints reliably indicate TF occupancy. For this purpose, we compared footprints from 21 DNase-seq experiments in lymphocytes to predicted TF binding sites (peak regions) from ChIP-seq of 66 TFs in the same cell type. The genomic background for this analysis is the set of all genome-wide instances of the sequence motifs recognized by a given TF. On their own, these motif instances have an extremely high false positive rate >90%. We used the footprints from all 21 samples to define two scores at each genomic location: (i) the “best footprint score”, defined as the highest score at this location in any of the samples; and (ii) the “footprint fraction”, defined as the proportion of independent samples with a non-zero footprint score. We then tested for a linear relationship between these footprint scores and the likelihood that a motif instance corresponded to a true-positive binding site from ChIP-seq, testing performance via the Matthews correlation coefficient (MCC), area under the receiver operator curve (AUROC), and area under the precision-recall curve (AUPR). The most accurate predictor was the best HINT20 score, which achieved a maximum MCC of 0.42, corresponding to AUROC > 0.9 (Fig. 2). The high AUROC was driven by true negatives, which comprise 3,936,242 of the 4,110,504 total observations. The vast majority of true negatives had low HINT scores. True positives often had a high HINT score, but high HINT scores also had a significant false positive rate (Fig. S3). It should be kept in mind that true and false positive here are “soft” assignments, since ChIP-seq experiments are themselves imperfect predictors of TF occupancy.

**Figure 2.**
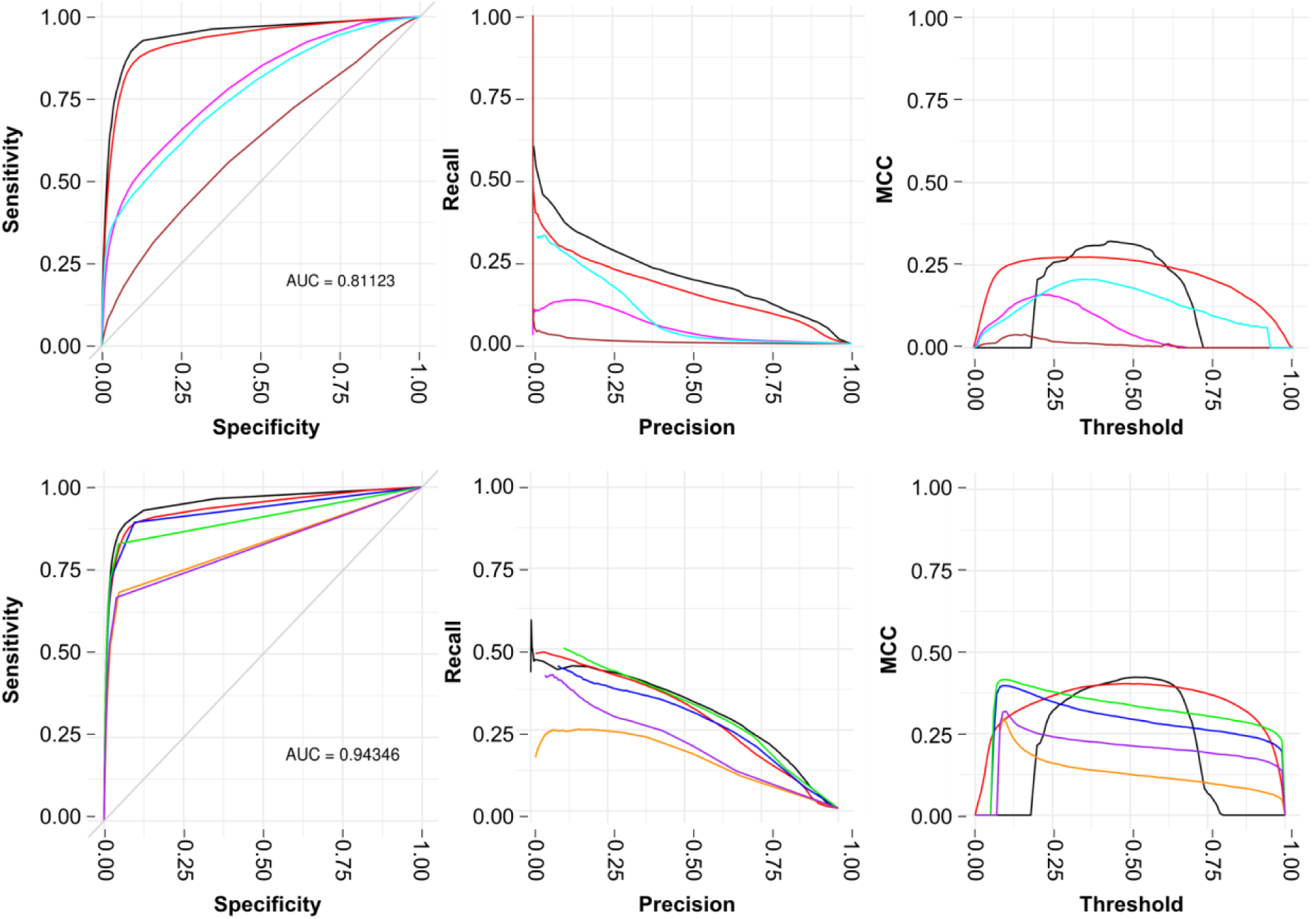
Predictive performance on a held out test set of a gradient boosted decision tree (GBDT) model of the 62 transcription factors (264 motifs) in the ENCODE-generated ChIP-seq samples. We compare to baseline models that use only motif information, TSS distance and GC content, and a linear model that uses all of these. A) Results using motifs devoid of footprint scores and metrics but including the following features: GC content, motif score, distance to TSS, and TF classes. B) Results for footprints generated from both Seed16 & Seed20 alignments using all aforementioned features, footprint scores and footprint metrics. The GBDT model obtains the best performance by nearly all metrics, though the amount by which it outperforms the linear model on the footprint data is in some cases marginal enough that an interpretable linear model may be preferred for some applications. The threshold in the third column refers to the decision boundary at which the continuous output of the models, which varies between zero and one, is thresholded and a classification decision is made. We note that the aggregate models obtain good performance over a relatively wide range of thresholds compared to the models using individual methods.

We were curious whether performance could be improved by combining footprint scores from multiple algorithms with additional information about genomic context. We employed a supervised machine-learning approach, treating the ChIP-seq peaks as true positives. We employed two machine-learning algorithms: linear regression and gradient boosting trees implemented with XGBoost. We constructed and evaluated a comprehensive model that included as predictors the footprint scores from both HINT and Wellington using both the 16 bp and 20 bp seed sizes. Additional predictors included a score for the strength of the match to the sequence motif, TF Class, GC content, and distance to a transcription start site (TSS). We compared this comprehensive model to predictions based on footprint scores alone, as well as to a baseline model that considered motif scores and genomic context but ignored footprinting data.

In the comprehensive model, gradient-boosting and linear regression achieved maximum MCC of 0.42 and 0.40, respectively (Fig. 2), The predictor with the largest contribution to accuracy was the best HINT20 score, followed by the HINT20 footprint fraction (Fig. 3). Prediction accuracy was lower in the baseline models, but remained better than chance (gradient-boosting, MCC = 0.32; linear regression, MCC = 0.27; Fig. 2). In these models, distance to the TSS was the most significant contributor to the prediction. While the maximum MCC of the HINT20 footprint-only vs. comprehensive models were identical (0.42), the footprint-only model had a relatively small threshold window within which both true positive and false negative error rates were well controlled. Therefore, incorporating information about genomic context does not dramatically improve prediction accuracy but could potentially improve the robustness of these predictions.

**Figure 3.**
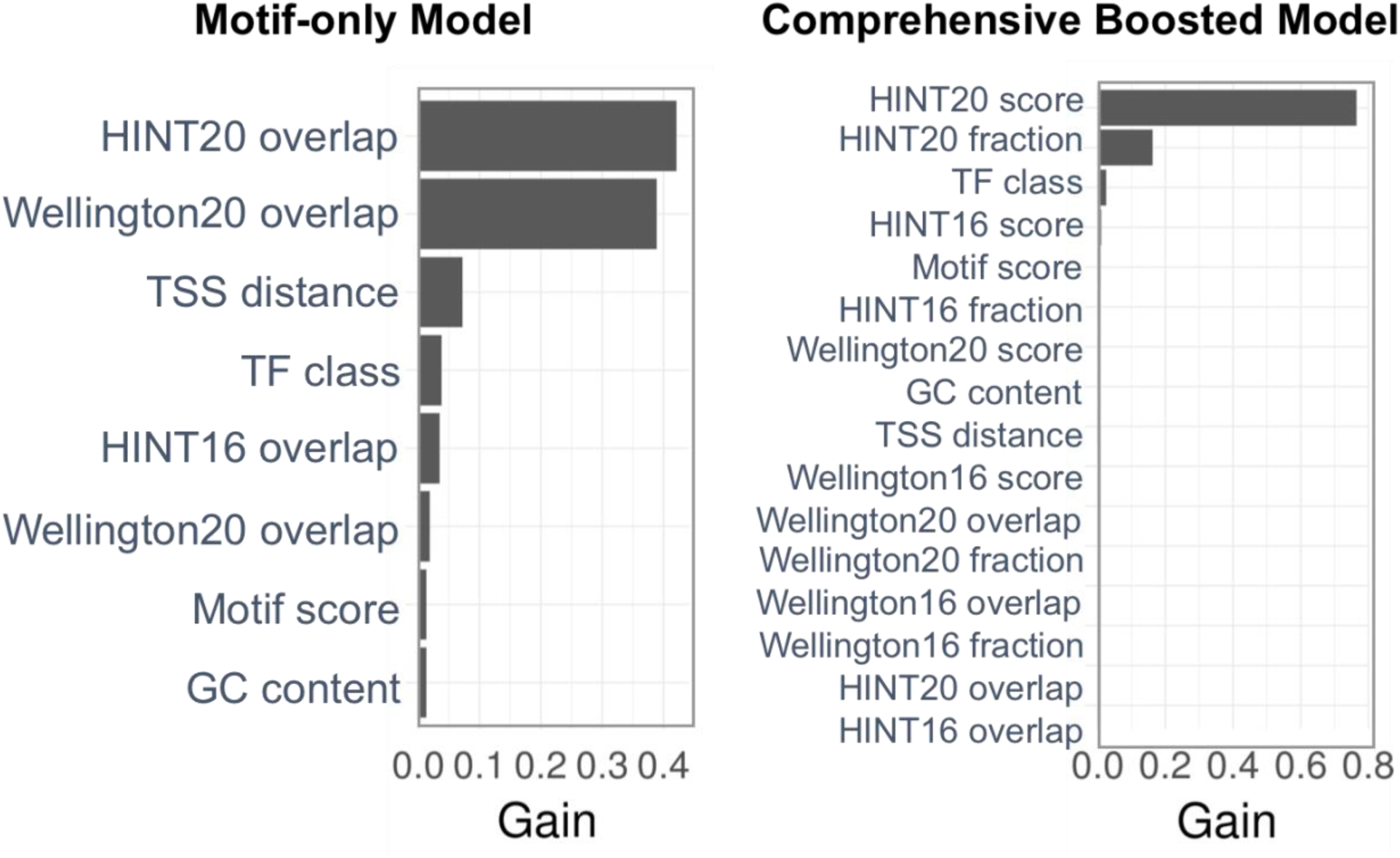
Importance matrix quantifying the contribution of each feature when trained and tested on the ENCODE ChIP-seq dataset for 62 transcription factors.

We used machine learning models to select appropriate cutoffs for high-quality footprints. We determined that a HINT score >200 and a Wellington scores < -27 were optimal filtering thresholds to control both false positive and false negative errors. Applying these filters reduced the percent coverage of the genome from 34% to 9.8% across all tissues (Figure 1B). This filtered estimate is in line with current estimates for the fraction of the genome that is actively involved in gene regulation. HINT20 footprints with scores >200 were used in downstream analyses unless otherwise specified.

### Footprints predict effects of genetic variants on gene expression

An important goal for footprinting is to predict the gene regulatory effects of non-coding SNPs. It has previously been shown that the majority of haplotypes with cis-acting effects on gene expression (expression quantitative trait loci; eQTLs) contain SNPs that are located within DNase I hypersensitive regions [32]. However, DHS regions span a large fraction of the genome, and many SNPs within DHS regions have no evidence for influencing gene expression. It remains controversial whether footprints more precisely capture the causal variants on eQTL haplotypes: some recent studies found that only a small fraction of eQTL haplotypes overlap footprints [8, 32], while others have suggested a stronger enrichment [33, 34]. To address this question, we examined overlap between footprints in our database with eQTLs from the Genotype Tissue Expression (GTEx) consortium.

We evaluated overlap between footprints (HINT20 score >= 200) from our database with expression quantitative trait loci (eQTLs) in 44 tissues from the Genotype-Tissue Expression Consortium (GTEx V6p) [12]. We focused on 1,561,655 genetic variants significantly associated with the expression of a nearby gene (<1MB) and in the credible set of variants for the gene at a high confidence level (95%) in at least one tissue, based on Bayesian fine-mapping with CAVIAR [35]. Across all eQTL and footprint tissues, we found that 163,330 of these 1,561,655 variants intersected a TFBS from our footprint database (TFBS-eQTLs). Counts of TFBS-eQTLs in individual tissues from our footprint database ranged from 743 (urinary bladder) to 71,692 (extra-embryonic structure) (Fig. 4A). We tested whether this overlap was greater than expected by chance by mapping footprints to all 11,959,406 genotyped and imputed variants in the GTEx V6p dataset, followed by resampling permutations. We found significant enrichments (p < 0.001) for all 27 footprint tissue x 44 eQTL tissue combinations. The overlap of footprints and eQTLs in mismatched tissues likely reflects the fact that many of the strongest footprints and eQTLs are detected in multiple tissues [12]. Sample size differs dramatically between tissues both in our footprint database and in GTEx, making it difficult to discern biologically relevant tissue-specific effects. Therefore, in subsequent analyses we considered all eQTLs together, regardless of the tissue in which they were discovered.

**Figure 4.**
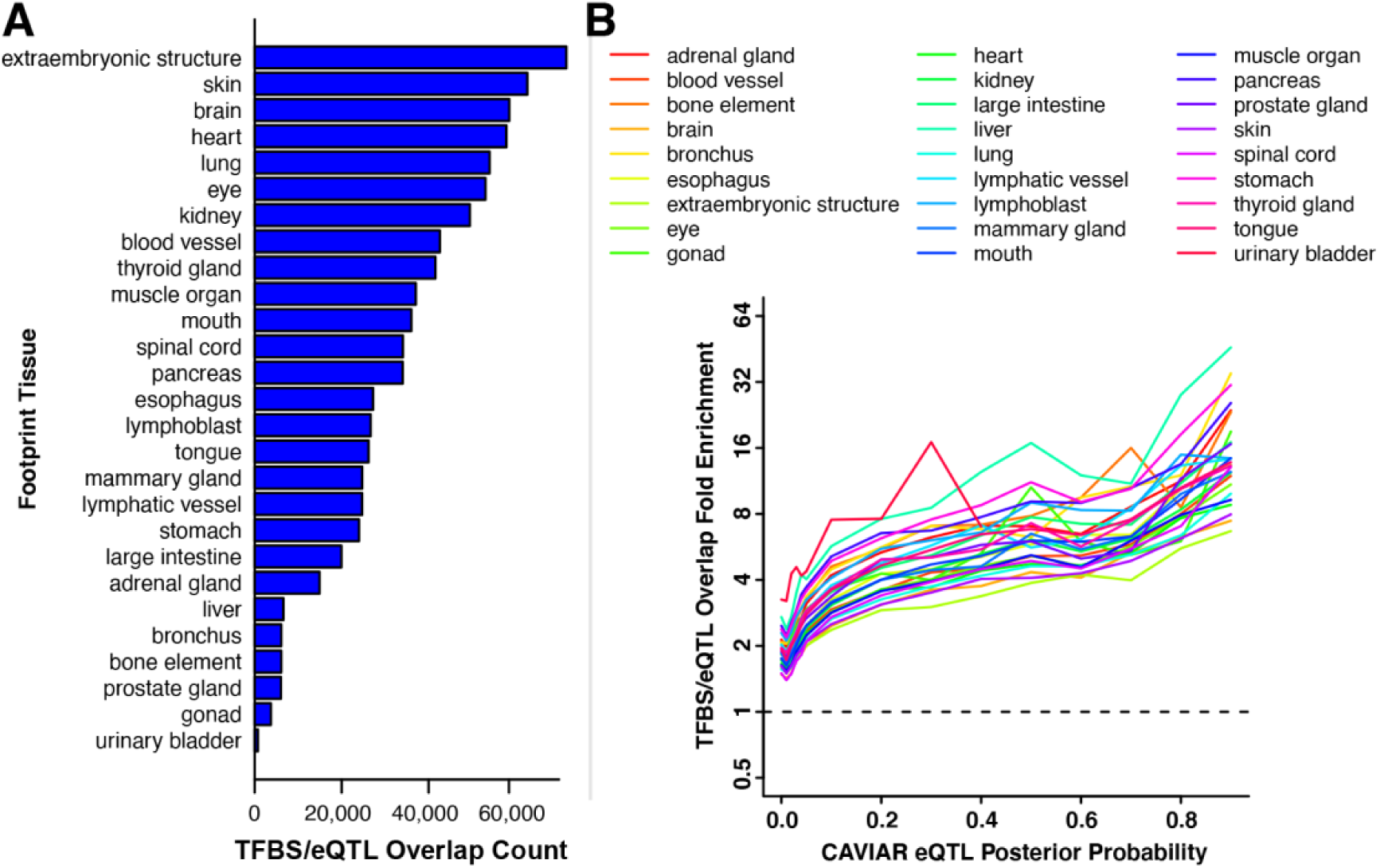
A) Counts of eSNPs overlapping predicted TF binding sites across all DHS tissues. Barplots indicate the total number of eSNPs overlapping footprints across all GTex tissues. B) Comparison of CAVIAR eQTL scores with the fold enrichment for TFBS/eQTLs.

We also determined whether eQTL SNPs with the highest likelihood of being causal variants from linkage-disequilibrium (LD)-based fine-mapping with CAVIAR were also the most likely to overlap a footprint. eQTL variants that overlapped a footprint had higher posterior probabilities for being causal than eQTL variants that did not overlap footprints (t = -61.4, p << 1e-308). Indeed, we detected a strong positive association between a variant’s posterior probability of being causal and the strength of enrichment for footprints, consistently across footprints from all 27 tissues (Fig. 4B). Focusing on the 3,193 eQTL variants with posterior probabilities > 0.8, we found that 29.2% (932) overlap a footprint. Resampling permutations indicated that this overlap for tissue-specific footprints is ∼10-40-fold greater than expected by chance. These results suggest that a large fraction of eQTLs may be explained by causal variants that alter TF binding sites, with many of these effects captured by footprints in our database.

### Tissue-specific footprints are enriched for disease-associated SNPs

Finally, we tested the hypothesis that high-scoring footprints are enriched for genetic variants associated with disease risk. To address this question, we studied genome-wide summary statistics from well-powered GWAS of eight immune-related traits and of 27 psychiatric, behavioral, and cognitive traits (Methods; Table S3). We hypothesized that heritability for immune traits would be specifically associated with footprints in lymphocytes, while heritability for neuropsychiatric traits would be specifically associated with footprints in the brain.

When considering all tissue-specific footprints from our database (HINT20 score > 0 in any sample), we found that footprints from brain tissue were strongly enriched for heritability for brain-related traits, and footprints from lymphoblasts were strongly enriched for heritability for immune-related traits. However, since the majority of base pairs that are open chromatin have a non-zero footprint score, this result is not distinguishable from previously reported enrichments of heritability in open chromatin. We therefore examined whether footprints with higher scores contributed more to heritability than footprints with lower scores. We used a partitioned heritability approach where we divided footprints into deciles based off of their maximum tissue-specific footprint scores. We found that footprints with the highest scores in brain contributed disproportionately to heritability to brain related traits but were not strongly associated with immune traits (Fig. 5A). Conversely, footprints with the highest scores in lymphoblasts contributed disproportionately and specifically to heritability in immune-related traits (Fig. 5B).

**Figure 5.**
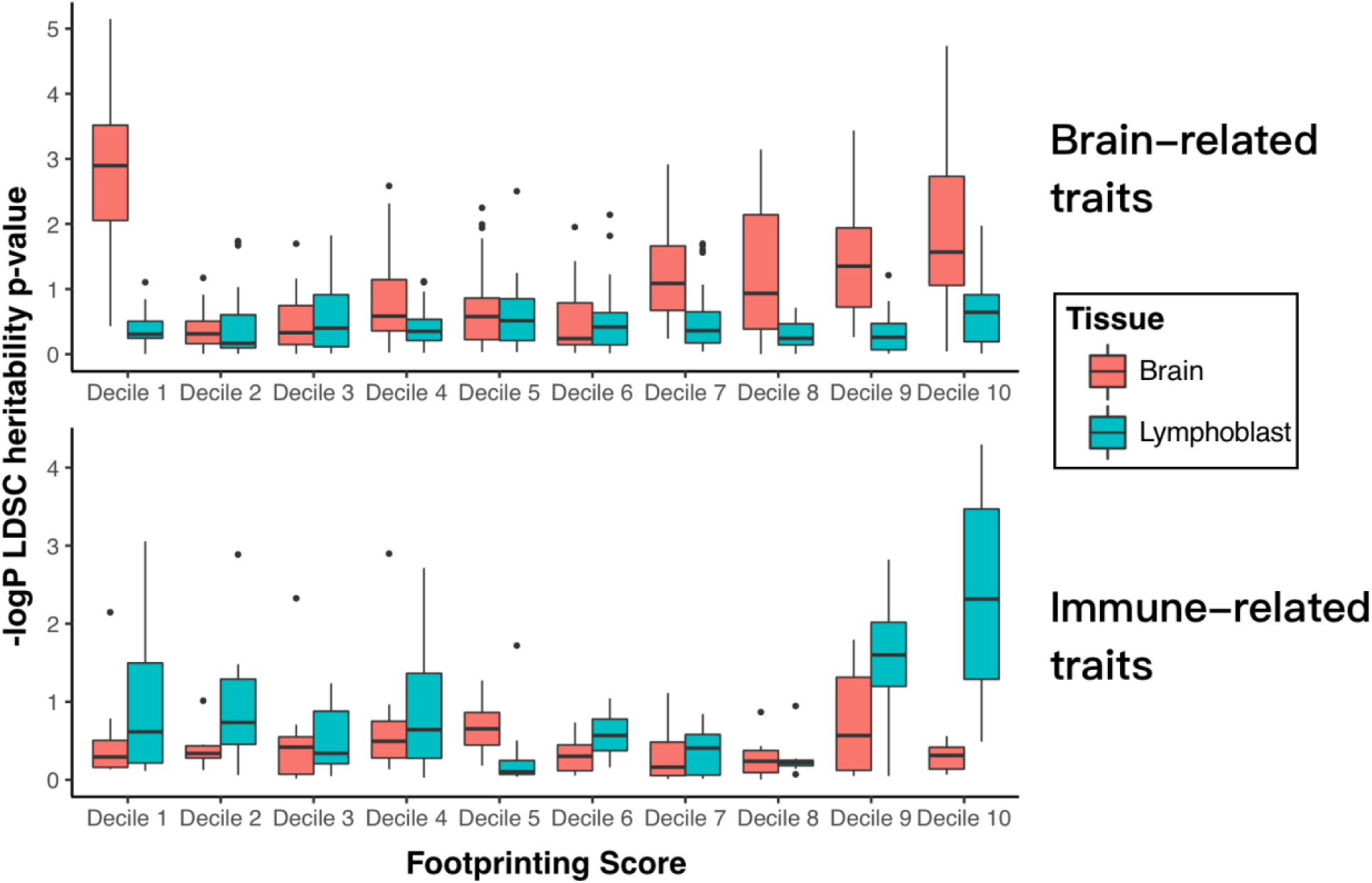
Partitioned heritability of tissue-specific footprints in related GWAS by footprint confidence score decile (decile 1 = lowest scores, decile 10 = highest scores). A) Partitioned heritability of brain footprints by decile in 27 summarized brain-related traits. B) Heritability of lymphoblast footprint deciles in 8 summarized immune-related traits.

Interestingly, we also found that positions of open chromatin in the brain that had very low footprint scores (bottom decile) contributed disproportionately to risk for brain-related traits. Motif enrichment analyses of the top vs. bottom deciles indicated that these segments of open chromatin are enriched for binding sites for distinct families of TFs. For instance, motifs recognized by several neurodevelopmental TFs (e.g., the LMX family) were disproportionately found in the bottom decile; these neurodevelopmental TFs are known to bind DNA more transiently than other TF classes, leaving a less distinct footprint signature [36] (Table S4). Taken together, our results support the hypothesis that the enrichment of disease risk in open chromatin is disproportionately due to variants that impact TF binding and indicate that a footprint’s score is positively associated with disease risk for many but not all TFs.

## DISCUSSION

Here, we have described a uniform workflow for DNase genomic footprinting and generated a comprehensive atlas of TF binding sites in 27 human tissues. We validated these footprints using data from ChIP-seq and eQTL experiments. At optimal thresholds, footprints in our database span 9.8% of the human genome, describing an expansive landscape of tissue-specific TF occupancy. We found strong, tissue-specific enrichments of footprints for disease-associated SNPs from GWAS, demonstrating the utility of our database to characterize gene regulatory mechanisms underlying human disease.

Machine learning approaches yielded several insights. First, footprinting information improved predictive accuracy compared to a baseline model. We note that since ChIP-seq itself is an imperfect gold standard, some footprints with no corresponding ChIP-seq may nonetheless be true binding sites for transcription factor. Footprinting may in fact identify a broader range of putative binding regions relevant to gene regulation, particularly in light of the strong relationship found with eQTLs. As a future direction, integration of additional epigenomic data could provide additional predictive power to discern active vs. inactive binding sites.

We also demonstrated strong enrichments of heritability for complex traits at the highest-scoring footprints, specifically in disease-relevant tissues. Given that the vast majority of risk variants in GWAS fall within non-coding regions, this finding suggests that disruption of TF binding may be a common mechanism by which genetic risk is conferred. These results build on previous findings that heritability for complex traits is enriched in open chromatin regions. Annotating risk variants with footprint scores improves specificity and mechanistic insight compared to annotating these SNPs based only on chromatin state. This finding demonstrates the utility of our footprint atlas for fine-mapping and other systems-level interrogations of complex genetic traits. Interestingly, we found that low-scoring footprints in the brain were highly associated with risk, and that these footprints disproportionately contained motifs for developmental TFs. This indicates that caution should be taken when using hard footprint score cutoffs, especially in the brain.

This resource also represents a case-study in the development of scalable cloud-based systems for large-scale data analysis [28]. The Globus Genomics [39] workflow used to create this resource can readily be extended to new open chromatin datasets and footprinting algorithms as they become available, potentially including newly developed approaches for open chromatin profiling in thousands of single-cells. This workflow is part of a family of interconnected tools being built within our Big Data for Discovery Science (BDDS) center (http://bd2k.ini.usc.edu). We have made user-friendly flat files for all footprints in this analysis available at http://data.nemoarchive.org/other/grant/sament/sament/footprint_atlas.

## ACKNOWLEDGEMENTS

This work was supported by the Big Data for Discovery Science Center of the NIH Big Data to Knowledge program (U54 EB020406; N.D.P. and L.H.), a contract from the National Institute of Aging (United States; U01 AG046139; N.D.P.), the NIGMS Center for Systems Biology at the Institute for Systems Biology (P50 GM076547; L.H. and N.D.P.), and by the national BRAIN Initiative (R24MH114788, Owen White, PI; R24MH114815, Owen White and Ronna Hertzano, PIs; 5R01HG009018: Hardening Globus Genomics. Ian Foster and Ravi Madduri, PIs).

## STAR Methods

### Lead Contact

Further information and requests for resources and reagents should be directed to and will be fulfilled by the Lead Contact, Seth Ament.

### Materials Availability

This study did not generate new unique reagents.

### Data and Code Availability

Footprint data files are freely available at http://data.nemoarchive.org/other/grant/sament/. Code and workflows available at https://github.com/globusgenomics/genomics-footprint.

### Key Resources Table

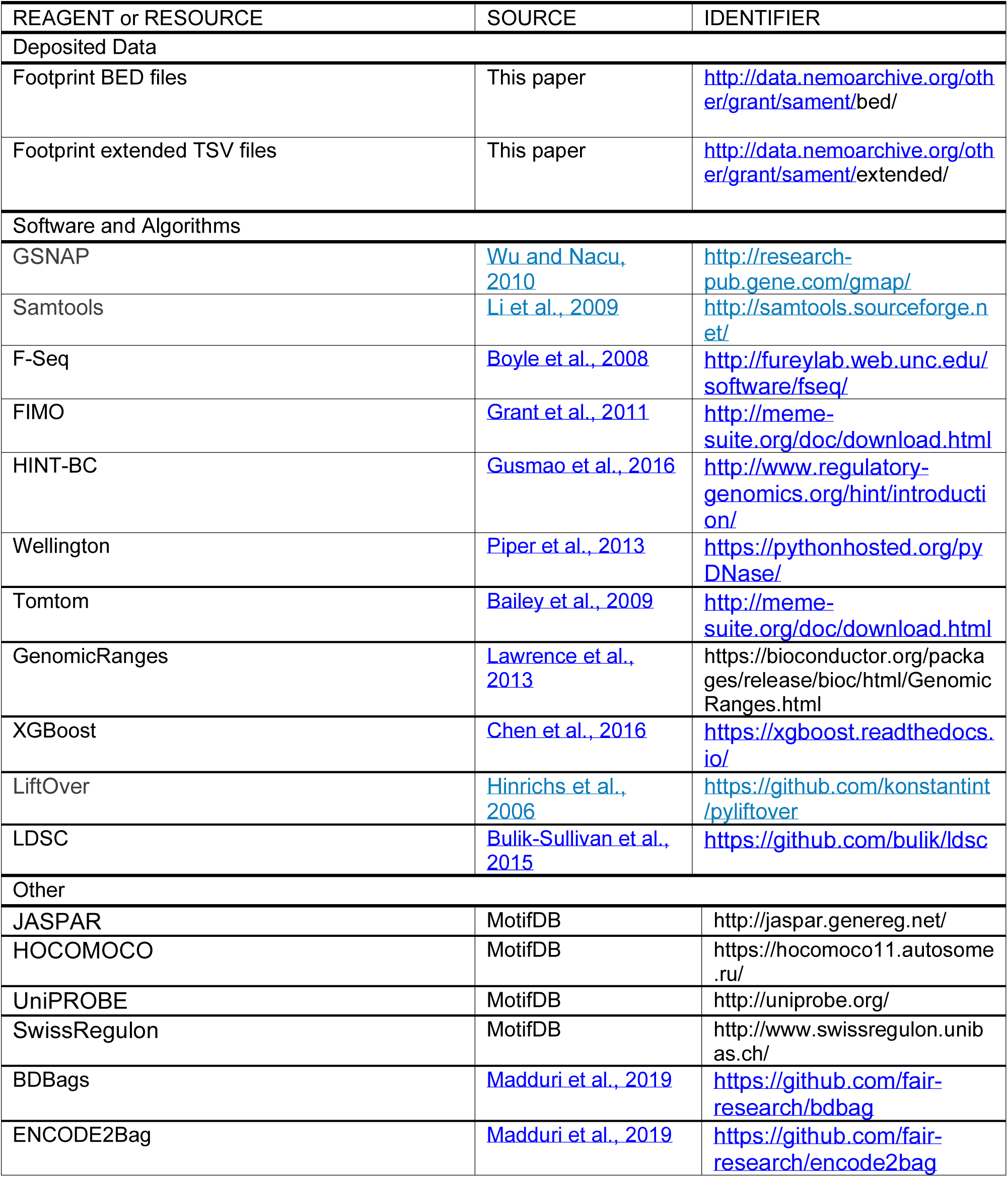
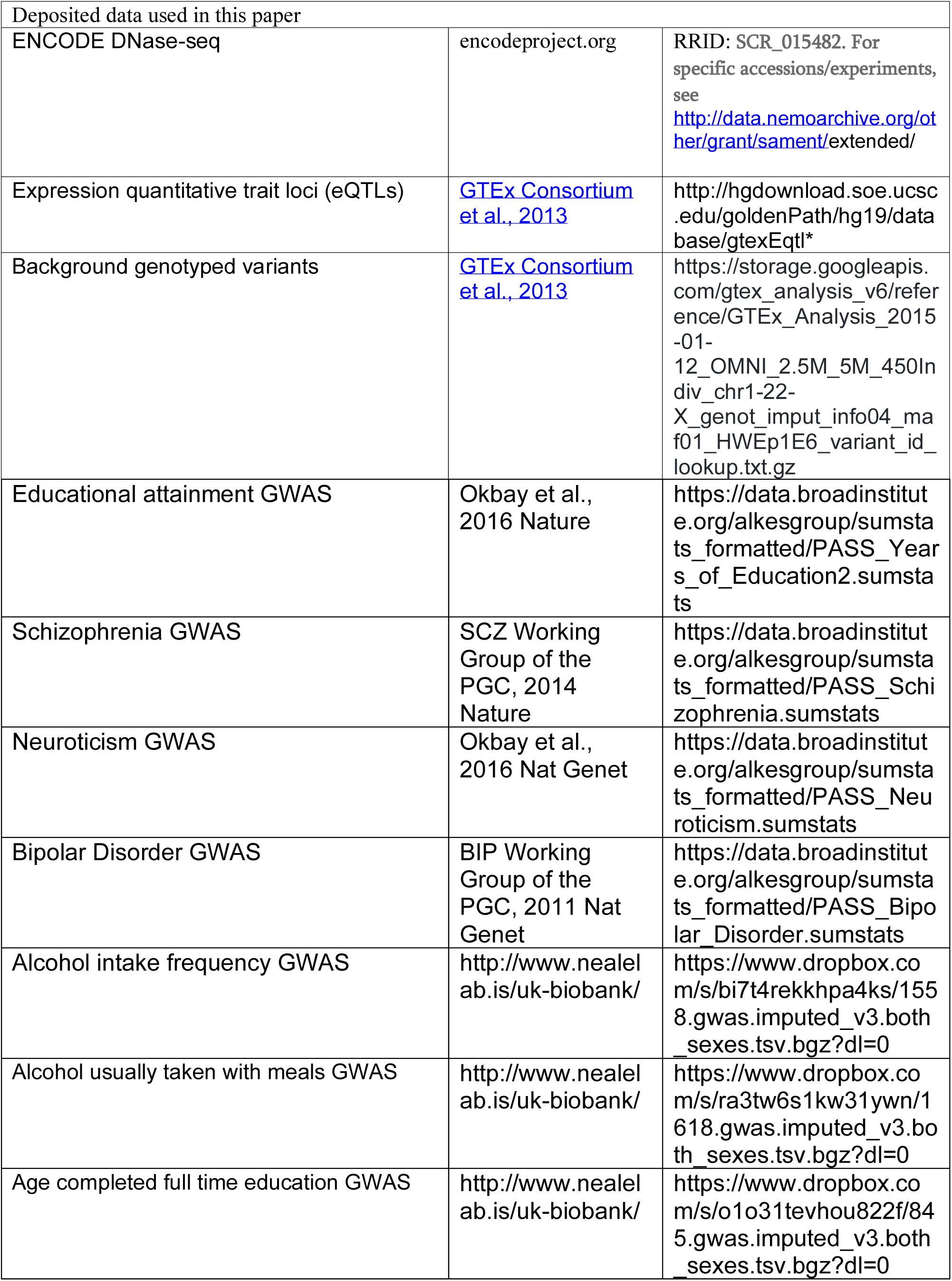
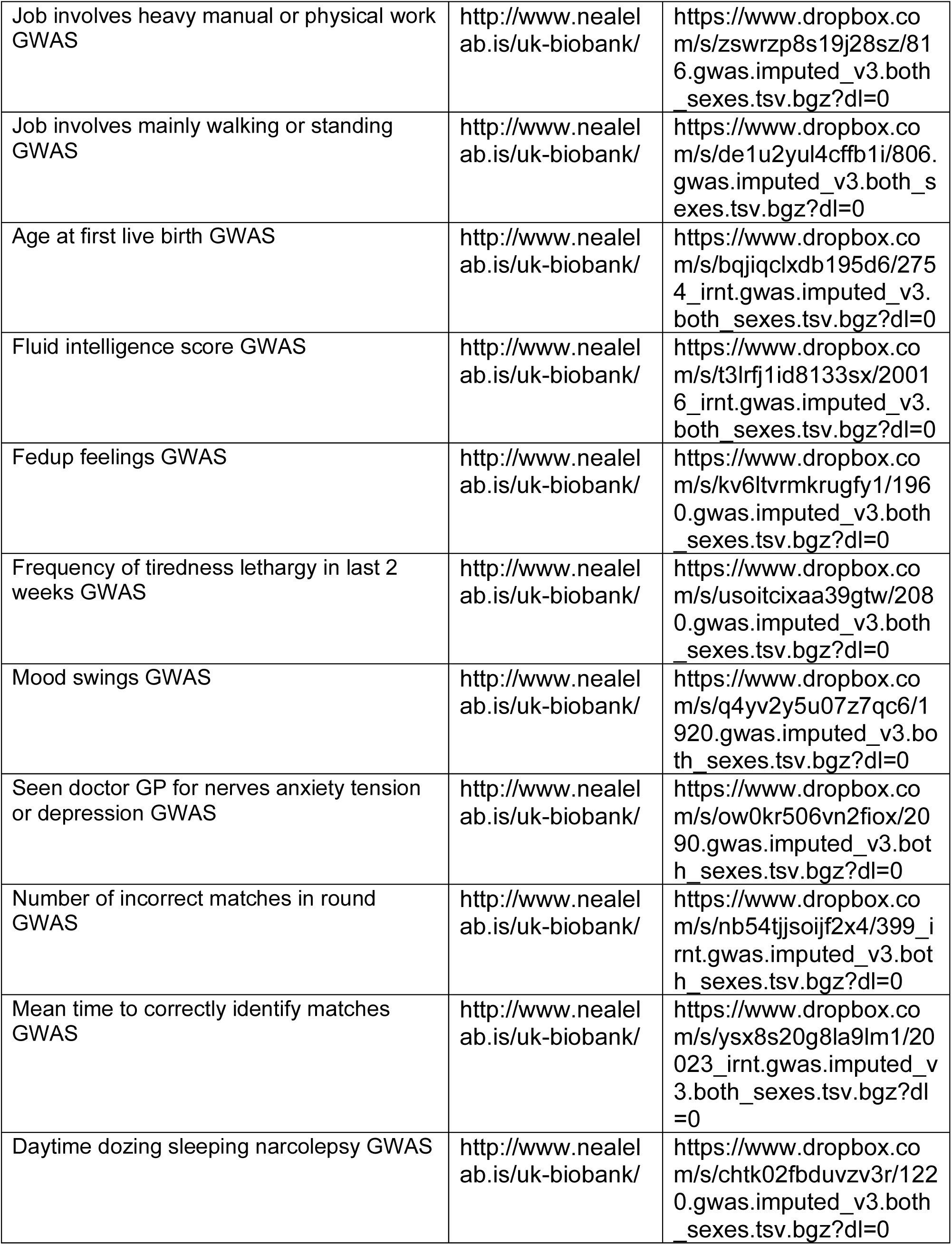
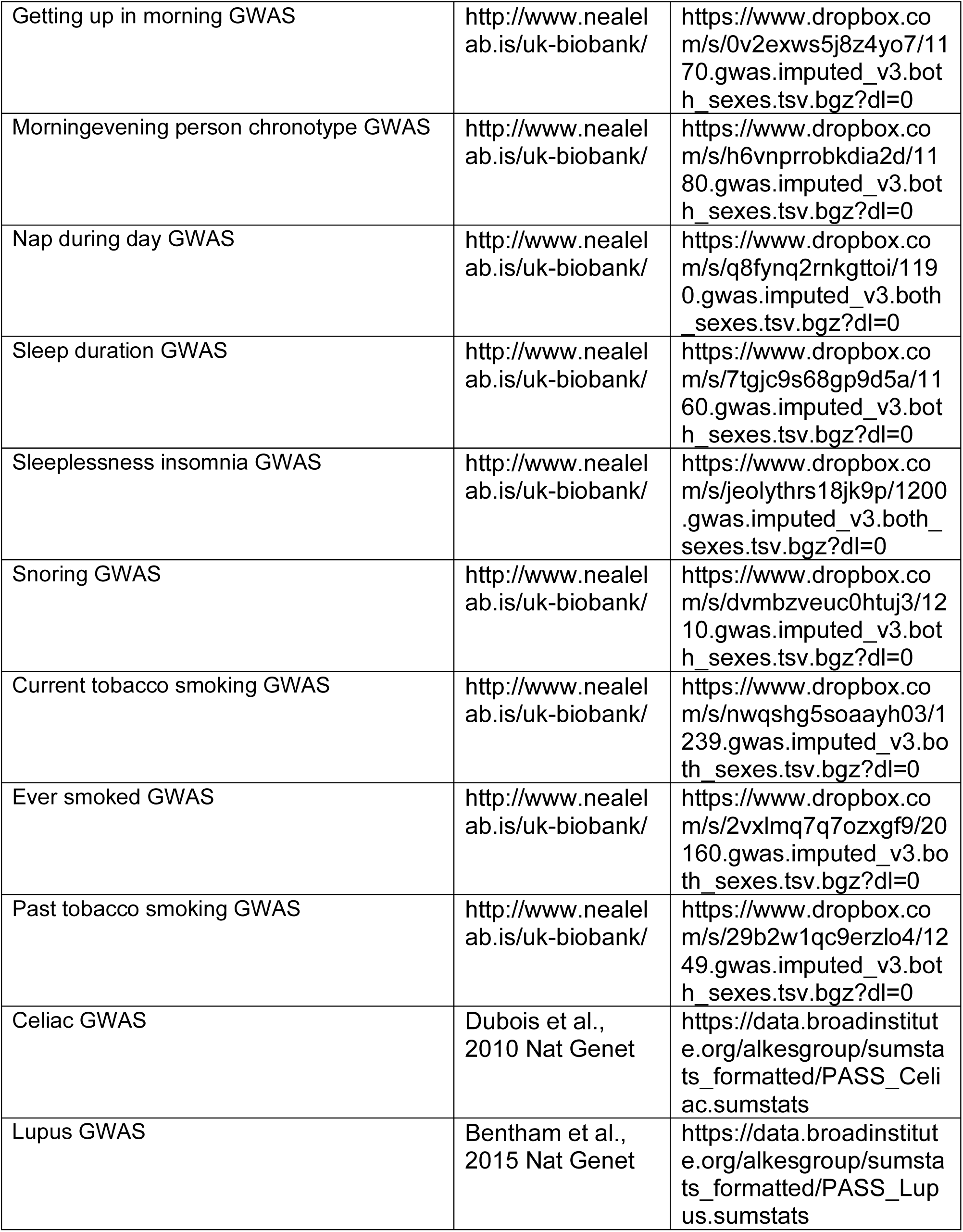
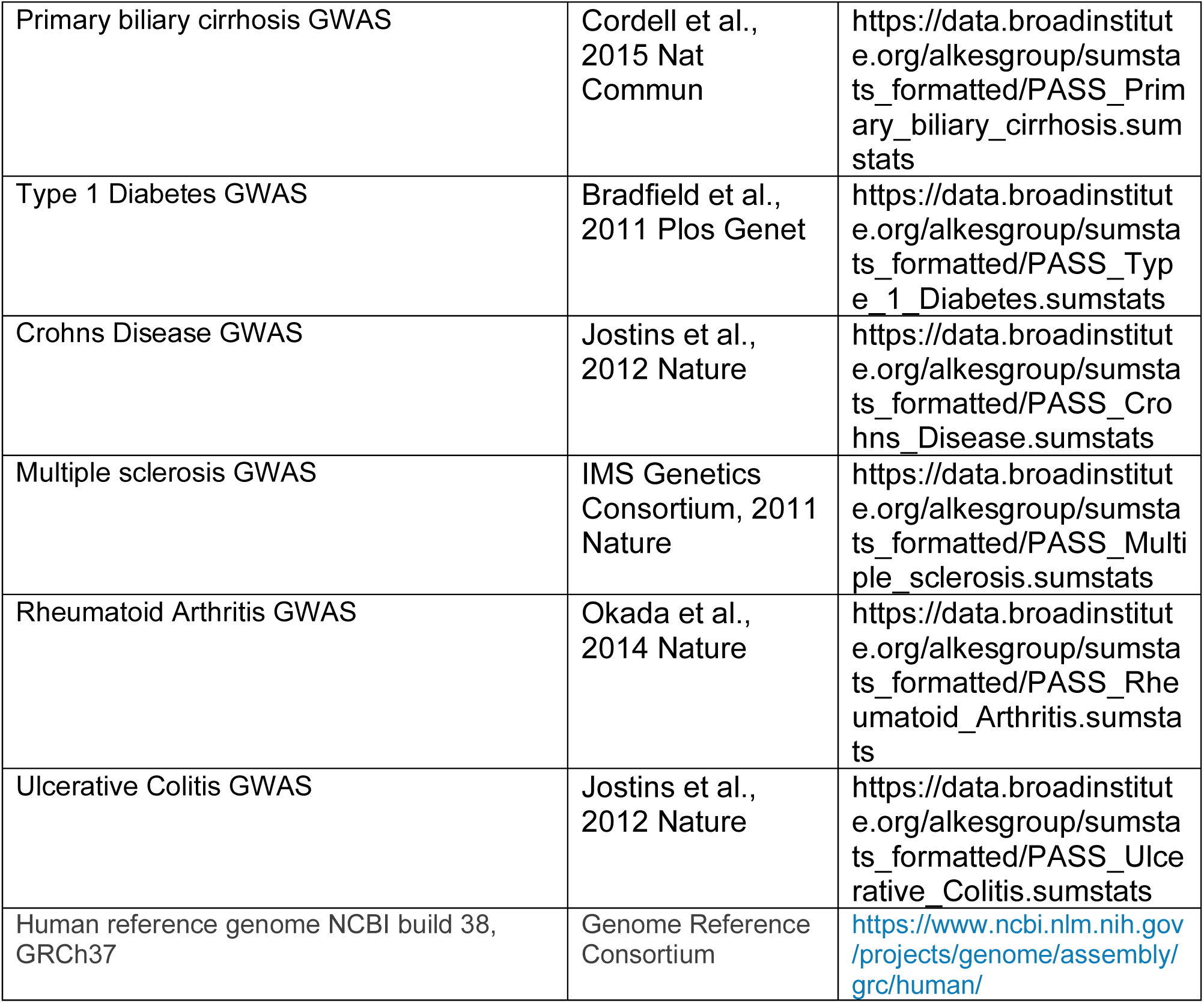

#### Overview

We created and executed footprinting workflows using various tools and services built and operated as a part of the NIH Big Data to Knowledge (BD2K) Big Data for Discovery Science (BDDS) center (http://bd2k.ini.usc.edu). At a high level, these tools enabled authoring and orchestration of complex and multi-tool workflows, transparent and elastic scaling on cloud resources, reproducible analysis based on provenance captured using minids and Big Data Bags (BDBags) (detailed below). The scalable workflows were built using the cloud-based Globus Genomics service [37]. These workflows include data retrieval from ENCODE using our ENCODE2Bag service that creates a portable data unit that encapsulates the entire results of an ENCODE query at a specific point in time. The resulting BDBag is passed as input to various analysis workflows that are executed in parallel to identify DNA footprints using cloud-based resources. The Globus Genomics platform, coupled with the BDDS tools, facilitates reproducibility of complex analysis for large cohorts through well-defined and published workflows [28].

#### BDBags, Minids

The input data from ENCODE consisted of all available DNase Hypersensitivity (DHS) datasets from 27 tissue types. ENCODE provides metadata for each tissue type which was exported and included in a BDBag [38]. BDBag is a format for defining a dataset and its contents by enumerating the data elements, regardless of their location, and for associating metadata. BDBags can be passed between services and materialized (by downloading data elements) only when needed. All datasets used in the workflow are identified using minids—a lightweight identifier for uniquely identifying a dataset. Minid and BDBag tools provide mechanisms for exchanging datasets by name, without regard for location or size, and with assurance that the data have not been modified.

#### ENCODE2Bag Service and Globus Genomics

The ENCODE2Bag service provides a simple web interface for exporting identified, verifiable collections of data from ENCODE. The service when given an ENCODE query, dynamically creates a BDBag that is stored on Amazon S3, and identified with a minid. The BDBag does not contain the large genomics files, but rather includes a manifest file which enumerates the files with their location(s) and checksum(s) for verifying integrity when accessed. The summary of the ENCODE query, represented as a Tab Separated Value file, is included in the BDBag as metadata to track and record provenance. Thus, given a BDBag, a user may, at any point in the future, obtain the results of that ENCODE query executed at the original time—an important property for reproducibility. BDBag tools abstracts the process by which a BDBag is “materialized”.

Globus Genomics is a cloud-hosted web service that enables rapid analysis of large genomics data. The service has over 3000 computationally optimized tools and a collection of best practices analysis workflows. Additionally, we added the data management tools built as part of the BDDS BD2K center to the service to make it easier for researchers to build high performance, reproducible bioinformatics workflows.

Globus provides reliable, secure, and high performance data transfer between Globus “endpoints” [39]. Globus provides a common interface to a variety of storage systems ranging from local POSIX file systems, through to cloud object stores (e.g., AmazonS3), high performance file systems, and even archival tape storage. Globus is able to orchestrate data transfer between any two systems by managing authentication with both endpoints, optimizing transfer configurations for transfer rate, recovering from errors, and notifying users of transfer status. We used Globus file transfer functionality to move large amounts of data from repositories, institutional storage systems, and local computers to the high performance, cloud-hosted compute resources used by the workflow.

The analysis workflows require only the minid of the input dataset to perform the analysis. The Globus Genomics service uses minid tools to transparently resolve the location of the BDBag, it then uses the BDBag tools to identify the contents of the dataset, and finally uses Globus to transfer the raw files to the cloud-hosted analysis infrastructure.

#### Scalable workflow for predicting Transcription Factor Binding Sites

In this workflow, we used the above-mentioned tools to materialize the BDBag for each tissue. Each tissue type contained DHS data for multiple samples. In addition, each sample had a variable number of replicate sequence data. Footprints were generated for the same input data using two alignment seed-lengths of 16 and 20 units, respectively. The analysis of the data consisted of aligning each replicate sample using the SNAP-aligner [29]. Once the alignment BAM files were produced for each replicate, they were merged using Samtools [40]. The merged BAM file was used to generate regions of open chromatin using F-Seq [16] based on the recommended parameters by Koohy et al, with the minimum reported size reduced from 500 bases to 400 [41]. Wellington was run with the -fdrlimit set to -1, to be the most lenient in reporting. HINT was run using standard settings. Neither Wellington nor HINT were run using any cleavage bias correction [17, 19]. The footprints were then stored in a relational database for ease of query.

The size of the input data (2.5 TB) and variability in replicate quantity for all samples (1591 FASTQ samples) made for a complex analysis (Figure 1). The Globus Genomics platform allowed us to automate this analysis through its support for transparent batch submission and parallelization methods. We utilized Amazon EC2 r3.8xlarge instance type with 32 CPUs and 244 gigabyte memory per node. The analysis of all tissues generated over 5 TB of data while using approximately 68,771 CPU hours (2149.1 node hours). The analysis of each tissue was executed in parallel. In addition, each patient and their replicates were executed in parallel, as well as each footprint algorithm

#### Alignment

For each tissue type, we started with the FASTQ files from the ENCODE portal (encodeproject.org). Some ENCODE experiments contain multiple biological samples, while others may contain only a single sample. An ENCODE experiment may contain single or paired-end reads, with varying depth of sequencing and varying read length in each experiment.

The ENCODE data was generated using short reads (<50 bases), resulting in a high number of potential sequence matches. This led us to produce alignments based on two different hash table seed lengths. Each FASTQ file (or paired-end files) was aligned to GRCh38 using the SNAP algorithm [29]. SNAP uses a default seed length of 20. We additionally aligned to seed size 16, given the shorter sequence lengths. Using the experiment groupings from ENCODE, we produced 386 BAM files for each seed.

#### Identifying regions of open chromatin

Based on work from Koohy *et al.*, who compared four different approaches (F-Seq, Hotspot, MACS and ZINBA) [41] we used F-seq [42] to identify regions of open chromatin from the aligned BAM files using the same recommended parameters. As stated in the F-Seq documentation, the results are non-deterministic because it uses a variable seed number in selecting a starting point for determining regions of open chromatin. The seed sets the sliding frame at which regions are considered, leading to slightly different beginning and ending points of open-chromatin. The resulting regions (in BED format) vary slightly when repeated. The variable coverage on the edges becomes less of an issue with increased sample numbers.

#### Motif database curation

As footprints from HINT and Wellington are motif agnostic and do not include information on motif matches, we integrated the footprint locations with motifs and motif-transcription factor mappings from JASPAR, HOCOMOCO, UniPROBE, and SwissRegulon. There is considerable redundancy between these databases, which often contain position weight matrices that are similar or identical. A motif in one database can also be quite different from the motif in another database associated with the same transcription factor, resulting in different mappings. To avoid inclusion of redundant motifs, we updated and modified an existing R package, MotifDB [43], to include the latest versions of all four databases. We evaluated the similarity of all motifs using Tomtom [44]. Those motifs that were significantly different from the 2016 release of JASPAR (-log(p-value) ≥ 7.3) were retained, yielding a total of 1,530 motifs. In addition to the mappings provided by each of the aforementioned databases, we also expanded the TF-motif mappings to incorporate families of TFs with very similar DNA sequence specificity, using information from TFClass [31]. The complete mapping can be accessed through MotifDB by calling the “associateTranscriptionFactors” method. The number of original motifs considered for each database and the number of motifs and transcription factor mappings retained after filtering are found in Supplementary Table 1.

Collectively, our aggregated collection of motif databases and mappings contains 1,530 unique motifs recognized by 1,515 transcription factors. Many motifs were associated with a single transcription factor, while a few promiscuous motifs were associated with as many as 60 transcription factors. Two representative examples of these mappings are found in Supplementary Figure 2. An entire map of all motifs and TFs can be found in the Supplementary Table 2. Reversing the association, many transcription factors were associated with one motif, while a few transcription factors were associated with > 100 motifs. The total number of motif-transcription factor mappings considered was 13,242.

#### Combining footprints with database of motifs

To maximize coverage, and because of the potential imprecise nature of footprints, if any part of a known motif overlapped with a single base of the footprint, an entry was created. Intersection was done by porting the motif instances and footprints into the GenomicRanges R package, using the “any” option.

### ChIP-seq validation and machine learning models

We joined all footprints based upon location in the genome to create one unified dataset per tissue. To account for the fact that the same footprints are often found in multiple samples from the same tissue, we retained the best score for each method and added as an additional metric the number of times a footprint was found at that location. As HINT is far more sensitive than Wellington, we scaled this count metric to one that captured the fraction of samples in which a given footprint was found. After we summed the number of footprints for each location, we used the highest number of occurrences as the denominator for all footprints in that method, resulting in a fractional representation for the occurrence metric. Additionally, we recognized that footprint-motif intersections include overlap of any size, but regions with higher overlap might indicate higher-confidence cases. To capture this effect, we calculated the overlap distance between each motif and its footprints for both seed as a fraction of motif length. JASPAR transcription factor class information was one-hot encoded in our feature matrix. GC content was calculated for each motif found within a footprint by using a window of 100 bases from the center on each side of the motif. Distance in base pairs (BP) to the nearest transcription start site (TSS) was calculated for each motif and transformed using the arcsinh (hyperbolic arcsine) function.

For purposes of validating the model, we designated chromosomes 2 and 4 as a hold-out set that was left untouched until the very end after all model parameter sets had been tested. Chromosomes 1, 3, and 5 were used to test the models as different parameters in architectures were explored. The remaining chromosomes were used to train the models. We trained two classes of models: 1) a basic logistic regression model, and 2) a gradient boosted model, which aggregates an ensemble of decision trees to learn a nonlinear decision boundary. Regression models were constructed for their ease of interpretability, as well as for a baseline to which we compare the performance of the boosted models. We trained logistic regression models not only for all features in the ensemble, but on each feature individually, in order to get an idea of which features were most predictive of ChIP-seq hits. The boosted model was chosen based on its predictive power, as gradient boosted trees have been shown to offer state of the art performance for tasks of this nature [45]. We used the R package XGBoost to create this model using a maximum tree depth of 7, 200 rounds of boosting, and a logistic regression optimization criterion [46].

One challenge that we encountered in creating this model is that the number of footprints for a given motif (or set of motifs connected to a given transcription factor) is orders of magnitude larger than the number of ChIP-seq peaks. This imbalance is problematic in the setting of this machine-learning format, as it increases memory requirements significantly and results in a poor signal-to-noise ratio. In order to address this issue in our training set, we sampled 20 million hits of 264 motifs, combined these motif hits with our lymphoblast footprints, then filtered for a 10:1 ratio of negative-to-positives. We did not filter any of the ChIP-seq hits in our training set. This resulted in a more balanced training set in which the features associated with true positives could be better learned. We also used a statistical measure of performance, the Matthews Correlation Coefficient (MCC), that was designed to be robust to unbalanced sample sizes in the two classes being compared [47].

### eQTL Enrichment

Expression quantitative trait loci (eQTLs) from the Genotype Tissue Expression Consortium (GTEx; V6p 95% credible causal sets) [12] were downloaded from the UCSC Genome Browser (http://hgdownload.soe.ucsc.edu/goldenPath/hg19/database/gtexEqtl+) on January 5, 2018. In addition, as a background set, we downloaded the table of all 11,959,406 genotyped and imputed variants from the GTEx V6p dataset (“GTEx_Analysis_2015-01-12_OMNI_2.5M_5M_450Indiv_chr1-22-X_genot_imput_info04_maf01_HWEp1E6_variant_id_lookup.txt.gz”) from the GTEx web portal (www.gtexportal.org; accessed March 16, 2018). GTEx variants were converted to hg38 coordinates using the UCSC Genome Browser’s liftOver tool (https://genome.ucsc.edu/cgi-bin/hgLiftOver) with default parameters. We identified TF binding site-altering variants by intersecting the locations of GTEx variants with the locations of TF binding sites from DNase-seq footprinting, using the genomic coordinates of motifs that overlap a footprint with a HINT score >= 200. Statistical associations between footprints and eQTL posterior probabilities were calculated using the t.test() function in R. Statistical significance for overlap between variants that alter TF binding sites and variants that are eQTLs was calculated from 1,000 re-sampling permutations, drawing variants at random from the complete set of genetic variants in GTEx V6p.

#### Partitioned Heritability Analysis

We utilized a partitioned heritability approach to characterize the relationship between footprint confidence scores and relevant phenotypes. First, we divided all the pooled footprints in a given tissue type into decile bins based on the score assigned to the best HINT20 score (1=lowest scores, 10=highest scores). We then used portioned LD Score Regression (LDSC) [25] to assess each decile’s contribution to heritability for several disease traits. The immune traits assessed were ulcerative colitis, type 1 diabetes, rheumatoid arthritis, primary biliary cirrhosis, multiple sclerosis, lupus, Crohn’s disease, and celiac disease. The neuropsychiatric traits included educational attainment, neuroticism, schizophrenia, and bipolar disorder, as well as 23 additional brain-related traits taken from the top 100 most heritable traits in the UK Biobank (Table S3) [48-57]. The top and bottom brain deciles were compared using a chi-squared test, and we used the residuals to determine over- and under-represented TFs in both deciles.

## Figure Legends

Supplementary Table 2. Motif-to-transcription factor mappings.

Supplementary Table 3. GWAS metadata from partitioned heritability analyses.

Supplementary Table 4. Chi-squared residuals for low-scoring footprints (Decile 1) vs. high-scoring footprints (Decile 9).

