## Supplemental Figures for "Atlas of Transcription Factor Binding Sites from ENCODE DNase Hypersensitivity Data Across 27 Tissue Types"

**Supplementary Table 1.** Motif database sources for FIMO matching. Motifs represents the number of motifs in the original database. Final Motifs represent the number of motifs used after running Tomtom. Total Mappings are the motif-to-transcription factor mappings found in the associated metadata and Total TFs are the number of transcription factors for which a mapping is found in the corresponding database.

| Origin | Motifs | Final Motifs | Total Mappings | Total TFs |
|---|---|---|---|---|
| Jaspar 2016 | 631 | 631 | 631 | 544 |
| HOCOMOCO v. 10 | 1066 | 649 | 1679 | 604 |
| UniPROBE | 380 | 162 | 1104 | 363 |
| SwissRegulon | 684 | 88 | 3835 | 684 |
| TFClass | NA | NA | 8570 | 762 |

**Supplementary Figure 1.** A) Footprints from the Seed16 alignment are compared to Seed20 alignment, considering the percent overlap. B) The number of footprints compared to the number of mapped reads for HINT and Wellington for all samples. C) The total number of hits (footprints and all intersecting motifs) per tissue type.

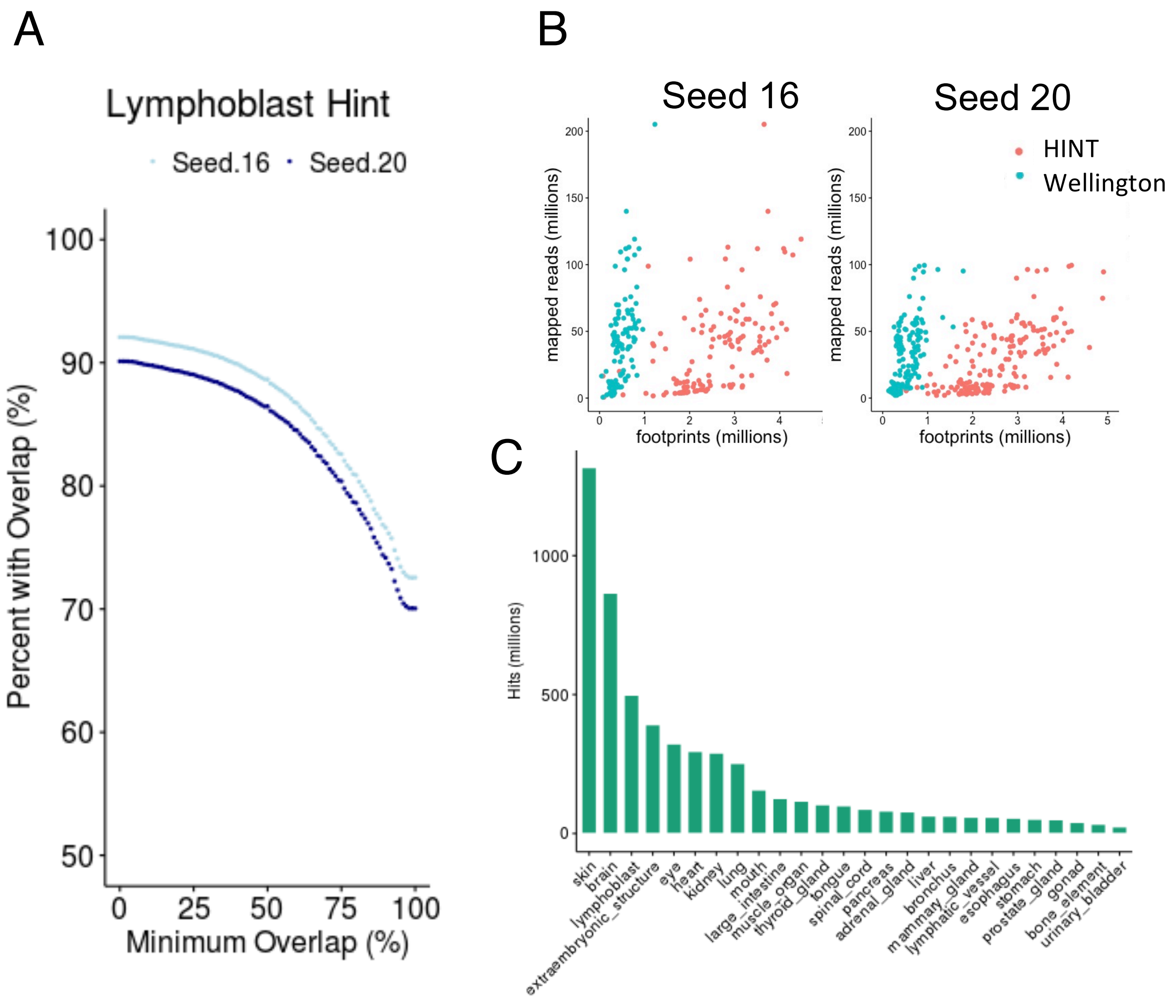

**Supplementary Figure 2.** Motifs from JASPAR, HOCOMOCO, UniProt and SwissRegulon were combined into a non-redundant set of 1,530 and matched to 1,515 TFs for a total possible combination of 13,242 pairings. Shown are representative sets of TFs and their corresponding motifs.

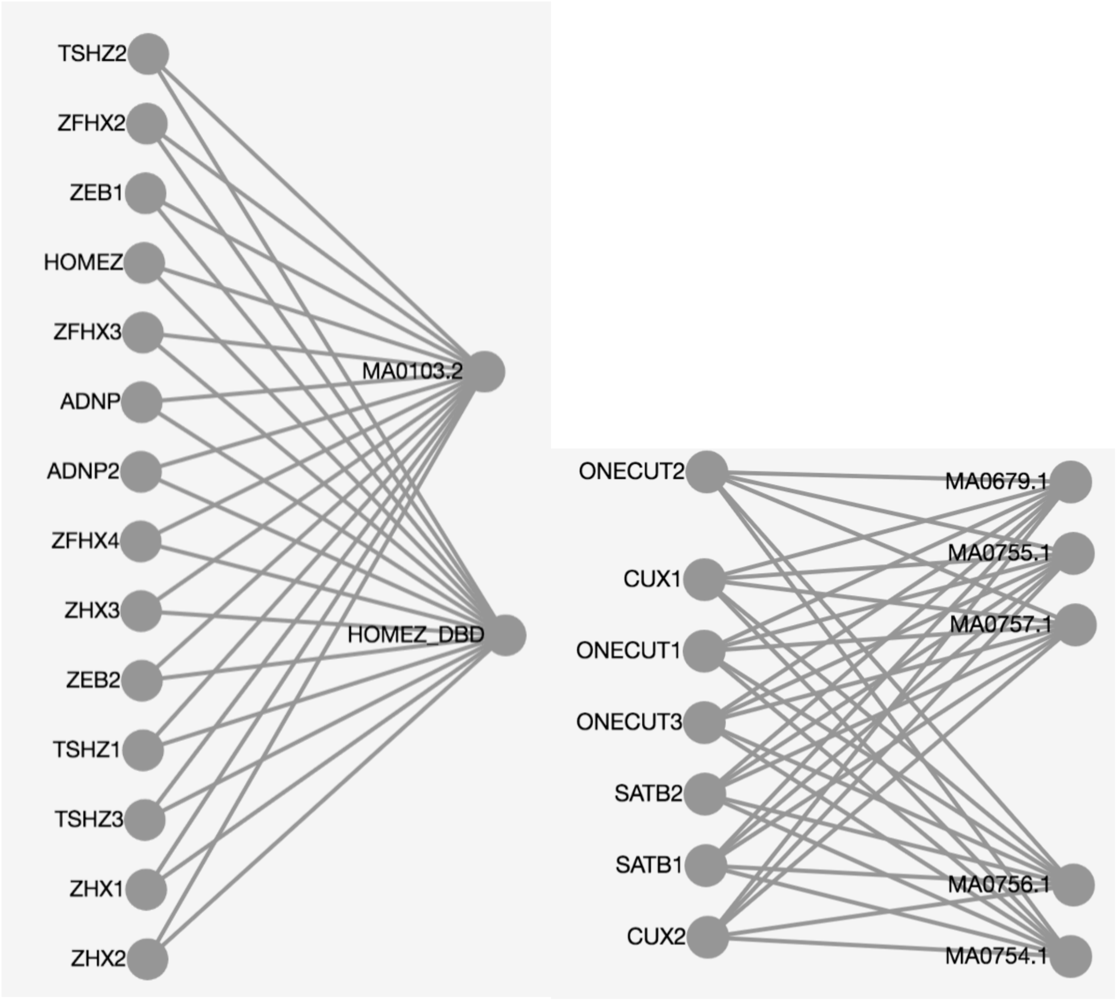

**Supplementary Figure 3.** Matthew's correlation coefficient from XGBoost model for all motifs associated with the transcription factor ELK1.

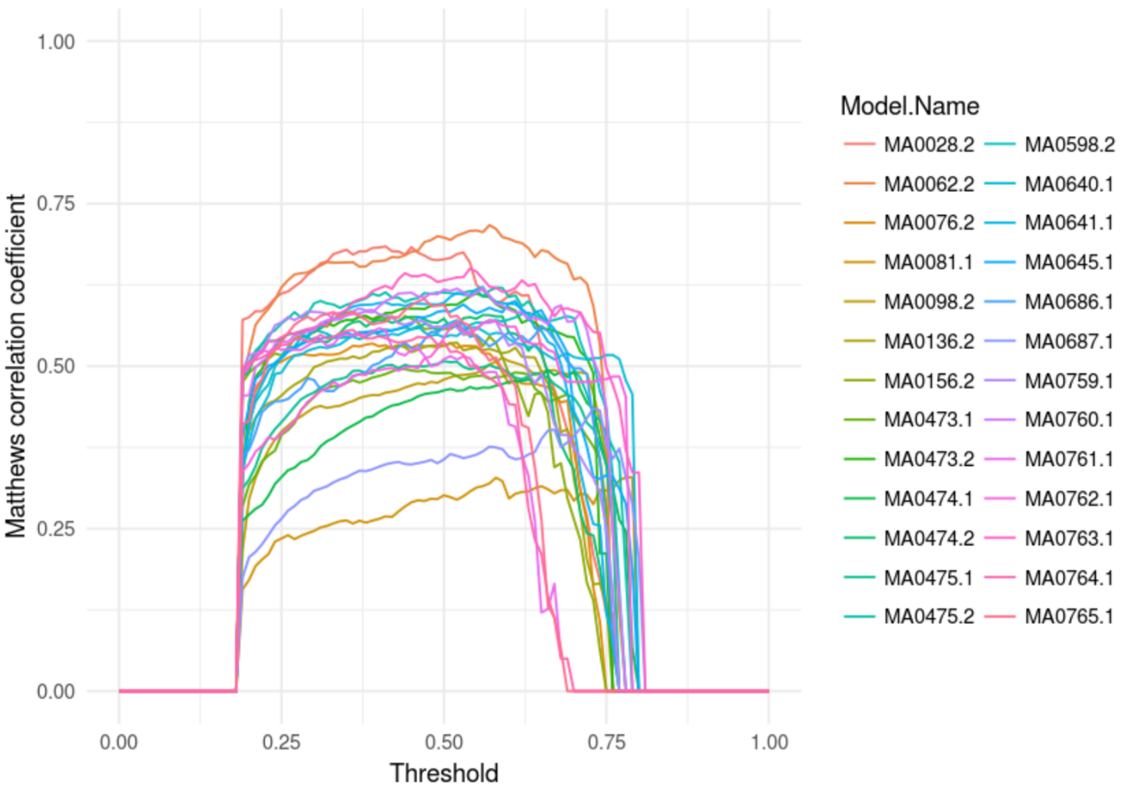

**Supplementary Figure 4.** The relationship between the predictions made by the machine learning is plotted on top of the two footprinting scores. The absolute value of the hyperbolic arc sine of Wellington 20 and HINT 20 score are on the x- and y-axis, respectively. The colors represent all possible prediction outcomes (false negative = 5,729, false positive = 140,952, true negative = 3,936,242, true positive = 27,579) from the boosted model.

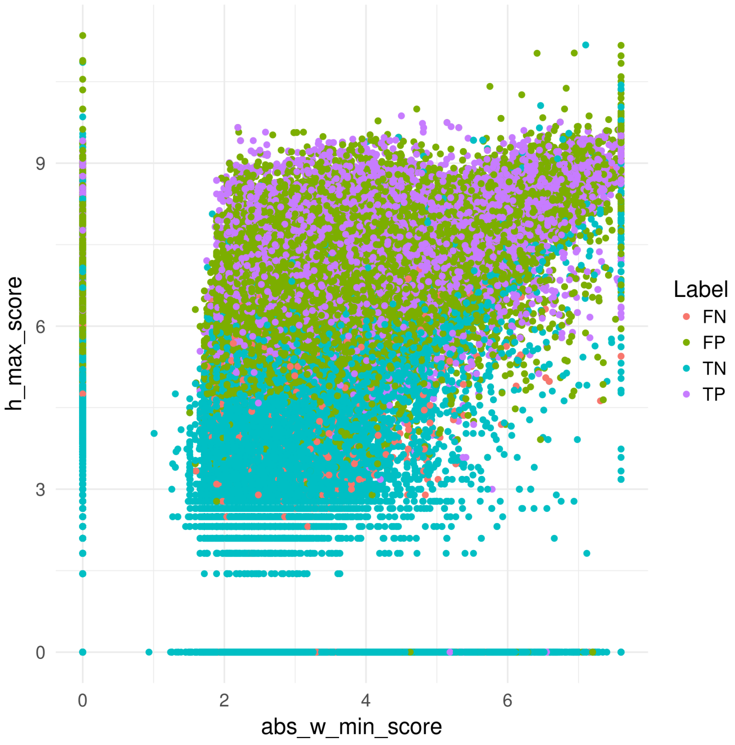
